# An origin of the immunogenicity of *in vitro* transcribed RNA

**DOI:** 10.1101/249714

**Authors:** Xin Mu, Emily Greenwald, Sadeem Ahmad, Sun Hur

## Abstract

The emergence of RNA-based therapeutics demands robust and economical methods to produce RNA with few byproducts from aberrant activity. While *in vitro* transcription using the bacteriophage T7 RNA polymerase is one such popular method, its transcripts are known to display an immune-stimulatory activity that is often undesirable and uncontrollable. We here showed that the immune-stimulatory activity of T7 transcript is contributed by its aberrant activity to initiate transcription from a promoter-less DNA end. This activity results in the production of an antisense RNA that is fully complementary to the intended sense RNA product, and consequently a long double-stranded RNA (dsRNA) that can robustly stimulate a cytosolic pattern recognition receptor, MDA5. This promoter-independent transcriptional activity of the T7 RNA polymerase was observed for a wide range of DNA sequences and lengths, but can be suppressed by altering the transcription reaction with modified nucleotides or by reducing the Mg concentration. The current work thus not only offers a previously unappreciated mechanism by which T7 transcripts stimulate the innate immune system, but also shows that the immune-stimulatory activity can be readily regulated.

## INTRODUCTION

Recent advances in RNA technology led to the emergence of RNA-based therapeutics (1–3). Promising therapeutic potential was shown for a wide range of RNAs, such as small interfering RNAs (siRNAs), aptamers, catalytic ribozymes, and mRNAs. Accordingly, there is an increasing demand for robust and cost-effective methods to prepare RNA on a large scale. While chemical synthesis can be utilized for relatively small RNAs, it is unsuitable for longer RNAs (>~20 nt) due to the exponentially decreasing yield with the increasing length of RNA. Enzymatic production using phage RNA polymerase, such as the T7 RNA polymerase (T7 pol), has been a popular method to prepare RNAs of various lengths. Advantages of the T7 pol include its robust activity and the ease of protein production. However, T7 pol is known to generate various kinds of aberrant byproducts, and its transcripts are often known to stimulate the vertebrate innate immune system (4,5). While the immunogenicity of T7 transcripts can be beneficial for certain applications (such as cancer immunotherapy), the current lack of understanding of the precise mechanism and the source of the immunogenicity limits harnessing such properties for therapeutic purposes.

RIG-I and MDA5 are two major cytosolic sensors that activate the innate immune system in response to viral dsRNAs (6). Upon viral dsRNA recognition, RIG-I and MDA5 activate antiviral signaling pathways that lead to the transcriptional up-regulation of the type I and III interferons (IFNs). Studies have shown that RIG-I and MDA5 have distinct RNA specificities, through which they recognize largely different groups of viruses (7–9). RIG-I recognizes dsRNA termini, in particular the 5’ triphosphate group (5’ppp), while MDA5 recognizes long (>~0.5-1 kb) dsRNA in a manner that depends on the duplex length, not 5’ppp (10,11). More detailed structural and biochemical analyses showed that MDA5 forms a filament along the length of dsRNA, and the filament formation is essential for the antiviral signal activation and dsRNA length detection (12,13). MDA5 also hydrolyzes ATP only upon binding to dsRNA, although ATP hydrolysis acts to regulate the MDA5 activity in seemingly complex ways (12).

It has long been thought that the immune-stimulatory activity of T7 pol transcripts is due to the fact that they harbor 5’ppp, which can stimulate RIG-I. However, data suggest that removal of 5’ppp by a phosphatase does not completely suppress the immunogenicity (14). Here we demonstrate that T7 pol often generates a high level of unintended double-stranded RNA (dsRNA) that are highly immune-stimulatory and made of the intended sense transcript and its fully complementary antisense transcript. The current work offers a previously unappreciated mechanism by which T7 transcripts stimulate the innate immune system, and a method to suppress such dsRNA byproduct formation.

## RESULTS

### *In Vitro* T7 transcription often generates dsRNA byproduct from a template designed for ssRNA

To examine the immune-stimulatory activity of T7 transcripts, we performed *in vitro* transcription of four independent RNAs, and tested their RIG-I/MDA5-stimulatory activities using the interferon-promoter driven dual luciferase assay in 293T cells. These transcripts were generated from templates that are designed to produce 512 nt ssRNAs with limited secondary structures (512A, 512B, 512C and 512D, see Table S1). Since 293T cells express a low level of RIG-I and little or no MDA5, we transiently expressed RIG-I or MDA5, stimulated the cells by transfecting RNAs of interest, and measured the signaling activity of RIG-I or MDA5 by the level of the luciferase activity. A known stimulator, polyinosinic-polycytidylic acid (polyIC), was used for comparison. Consistent with previous reports about the immune-stimulatory activity of T7 transcripts, we observed that all four T7 transcripts robustly stimulated both RIG-I and MDA5 (Figure 1A). Treatment of these RNAs with Calf Intestinal Phosphatase (CIP), which removes 5’ppp, largely suppressed the RIG-I stimulatory activities, but not the MDA5-stimulatory activities (Figure 1A).

**Table S1.**
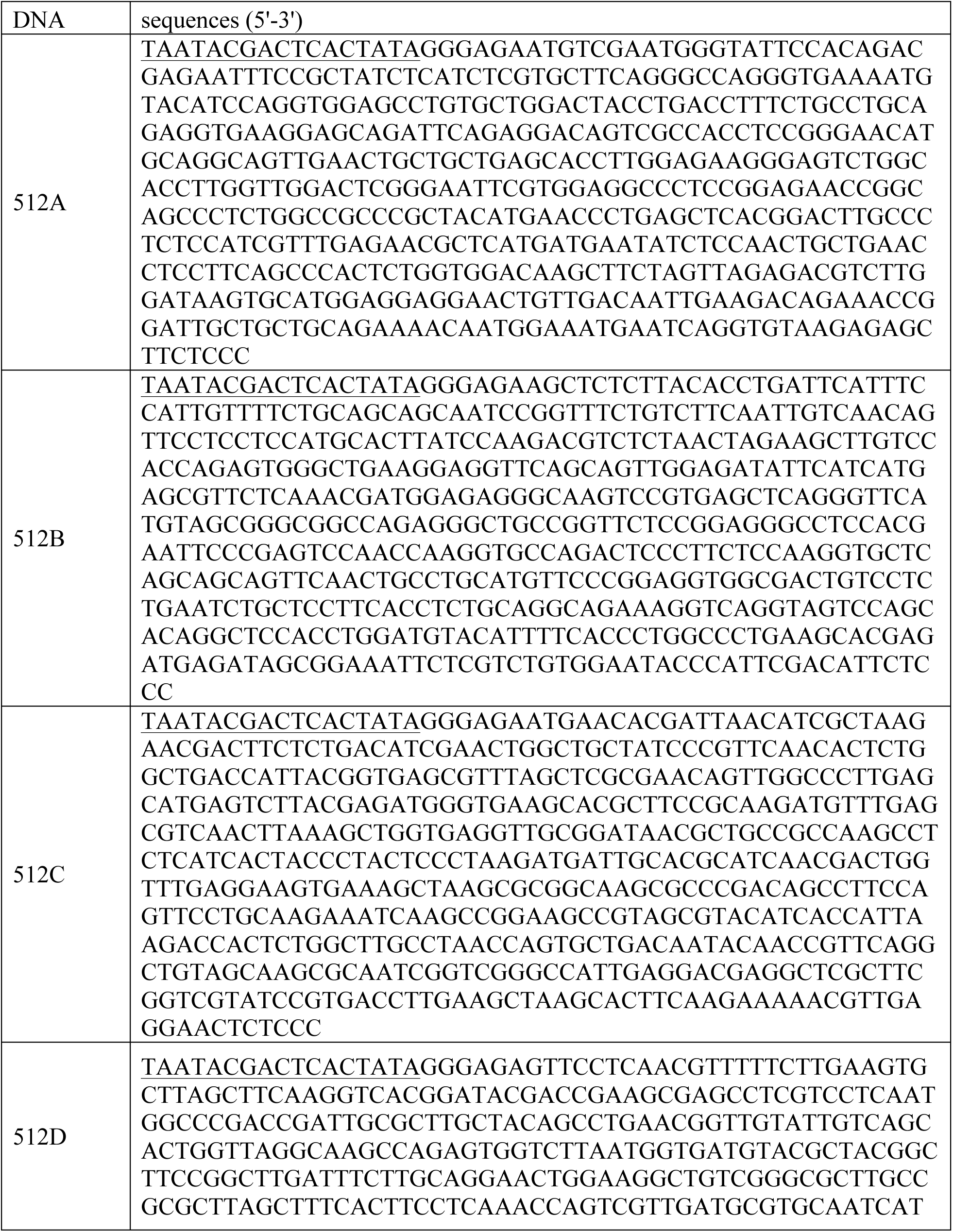
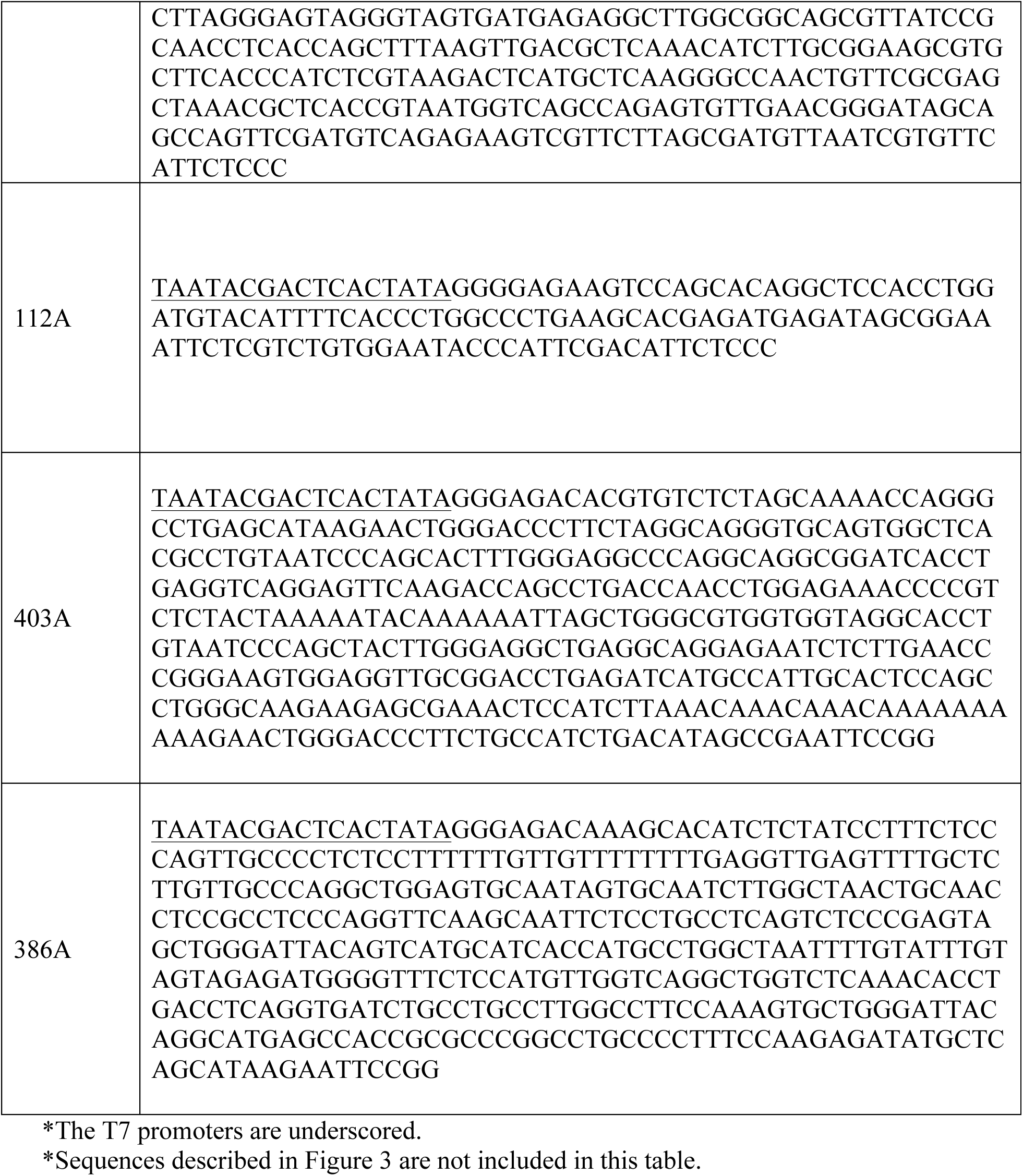
Sequences of the DNA templates used in this study*

**Figure 1.**
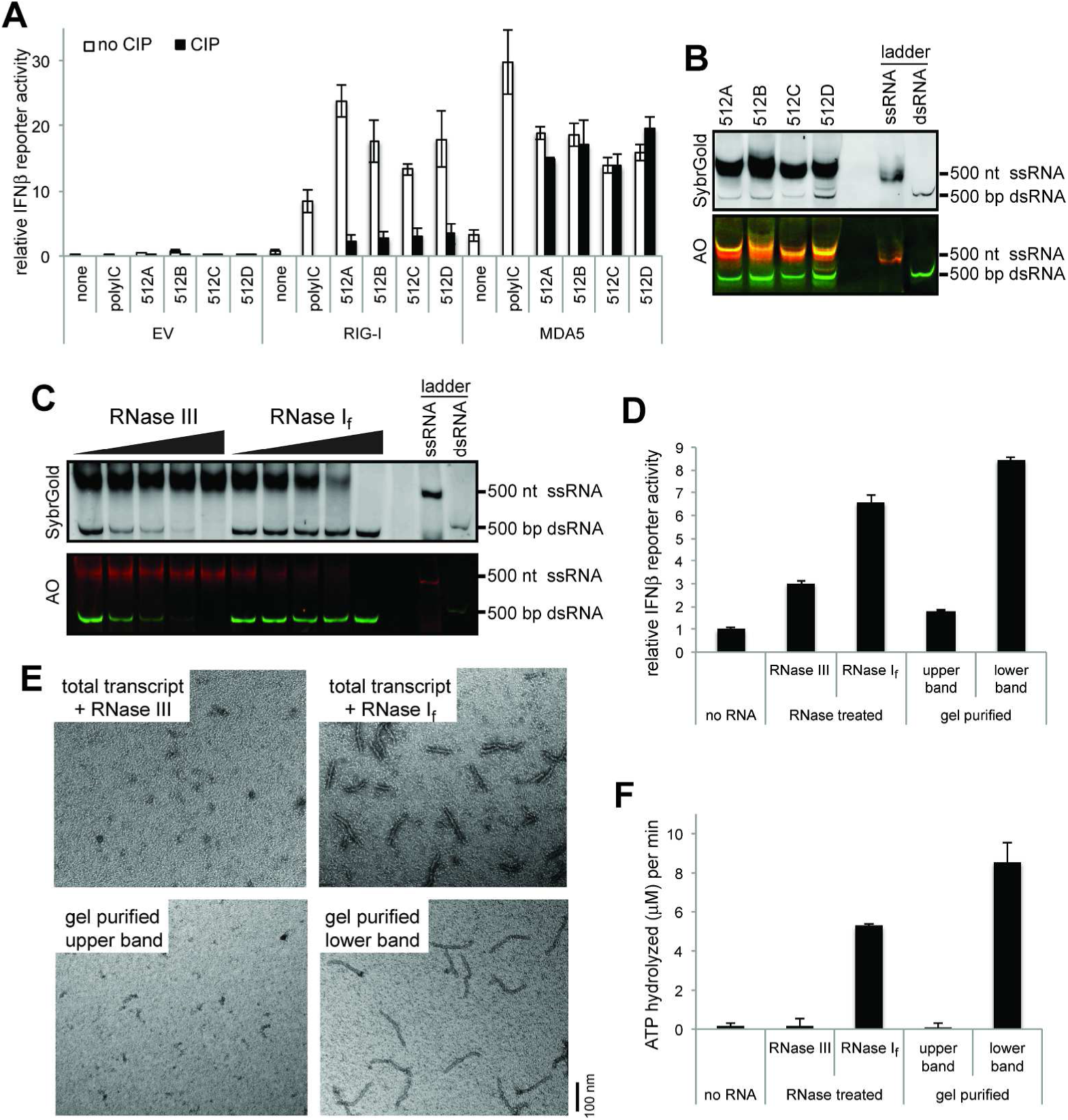
T7 pol generates MDA5-stimulatory dsRNAs from templates designed to encode ssRNAs. A. IFNβ promoter-driven dual luciferase assay. 293T cells were transiently transfected with the plasmids expressing human RIG-I or MDA5 (or empty vector control, EV), and stimulated with T7 transcripts for 512A, 512B, 512C and 512D. The RIG-I− and MDA5-stimulatory activities were measured by the dual luciferase activity. The stimulatory activity was compared before and after CIP treatment to examine the effect of 5’ppp. B. Native PAGE analysis of T7 transcripts for 512A, 512B, 512C and 512D. RNAs were visualized by SybrGold staining (upper panel) and Acridine Orange (AO) staining (lower panel). The 500 nt ssRNA and 500 bp dsRNA markers were used as controls. C. RNase-susceptibility analysis. The T7 transcript for 512B was treated with an increasing concentration of RNase I_f_ (1/40, 1/20, 1/14 and 1/6 units/l) or RNase III (1/4500, 1/3000,1/2000, and 1/1000 units/l), and was analyzed by native PAGE. D. MDA5-stimulatory activity of upper and lower bands of the transcript for 512B, as measured by IFNβ promoter-driven dual luciferase assay. The upper and lower bands were isolated by either RNase I_f_/III digestion or gel purification, and their MDA5-stimulatory activities were measured as in (A). All RNAs were treated with CIP to suppress the endogenous RIG-I activity. E-F. MDA5-stimulatory activity of upper and lower bands of 512B, as measured by MDA5 filament formation (E) and ATP hydrolysis (F). Both assays were performed using the RNA binding domain (MDA5∆N), which can form filaments and hydrolyze ATP as the full-length protein. Filament formation was detected using 2 ng/ul of RNA and 450 nM MDA5 by negative stain electron microscopy (EM). The ATPase activity was measured using 2 ng/ul of RNA and 500 nM MDA5.

To identify the origin of 5’ppp-independent MDA5-stimulatory activity, we analyzed the T7 transcripts by non-denaturing PAGE. Two bands were observed for all four RNAs tested, with the upper bands at an expected position near the 500 nt ssRNA marker, and the lower bands at a position near the 500 bp dsRNA marker (Figure 1B). While there were batch-to-batch variations in the relative levels of the two bands, they were the most prominent bands in all cases. The two bands displayed different fluorescence of acridine orange, a dye that binds RNA regardless of its secondary structure, but fluoresces differently depending on the RNA secondary structure (Figure 1B). The upper band showed a similar fluorescent behavior as the 500 nt ssRNA marker (rendered orange in the gel image), while the lower band did as the 500 bp dsRNA marker (rendered green). The two bands also displayed different sensitivities to RNase I_f_ and RNase III that are specific to ssRNA and dsRNA, respectively (Figure 1C, only shown for 512B as a representative example). The upper band was susceptible to RNase I_f_, while the lower band was to RNase III. Consistent with this notion that the upper band is mostly single stranded, while the lower band has significant duplex structures, only the lower band RNA, as purified by the gel extraction or RNase I_f_ digestion, had a significant stimulatory activity for MDA5-mediated antiviral signaling (Figure 1D). *In vitro* analysis of the interaction between the upper or lower band RNAs of 512B and MDA5 also showed that only the lower band can stimulate MDA5 filament formation and its ATPase activity (Figures 1E & 1F). Intriguingly, electron microscopy analysis of the MDA5 filament showed that the lower band RNA supported formation of ~150-160 nm long MDA5 filament (Figure 1E), a length expected for ~500 bp dsRNA (15). This suggested that the lower band is not the intended ssRNA with an intrinsic secondary structure, but is instead a ~500 bp dsRNA byproduct.

### dsRNA byproduct results from antisense transcription from the promoter-less DNA end

Previous studies reported that T7 pol can generate erroneous products by extending the 3’ end of the RNA with the sequence complementary to the intended RNA product (16). Such byproducts would fold onto themselves and form a hairpin structure with its duplex size equivalent to that of the intended ssRNA. To examine whether the ~500 bp dsRNA byproduct is produced in this fashion, we examined the 3’ end sequence of the intended ssRNA transcript (512B) using 3’ -RACE followed by Sanger sequencing (Figure 2A). The sequencing result shows that the ends are well-defined although a few nucleotide heterogeneities were also observed, a well-known behavior of T7 pol. This result is inconsistent with the idea that a long complementary 3’ extension is responsible for the observed ~500 bp duplex byproduct.

**Figure 2.**
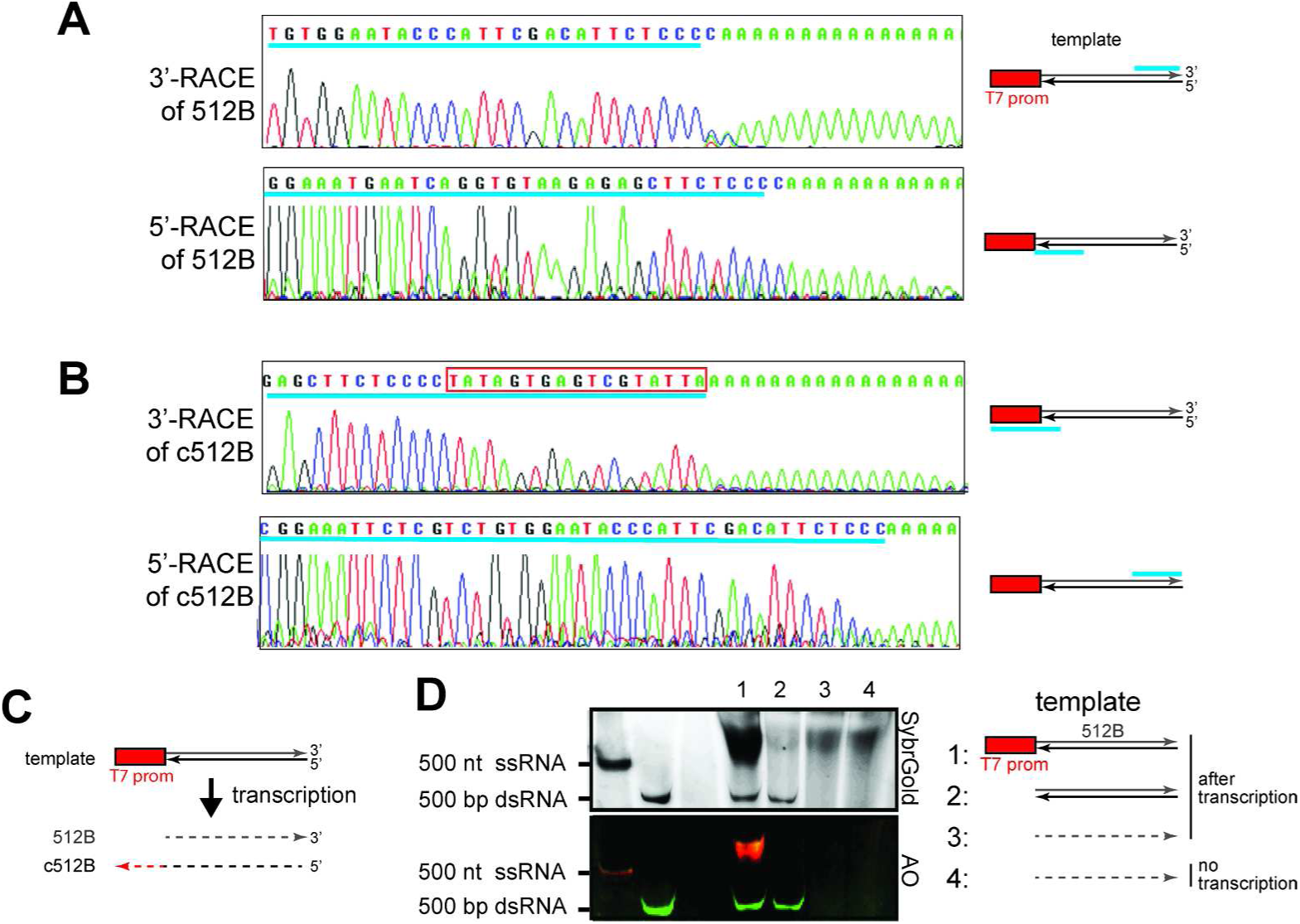
The dsRNA byproduct is formed by sense and antisense RNAs generated in promoter-dependent and −independent manners, respectively. A-B. Transcriptional start sites and end sites for the intended 512B product (A) and its complementary RNA byproduct (c512B) (B), as examined by 5’- and 3’-RACE. Transcriptional start and end sites are shown upstream of the poly A tail in the 5’- and 3-- RACE sequences, respectively. Cyan underscores in the sequence chromatograms indicate sequences matching those in the template. The location of the matching sequence in the template is shown in the schematic on the right. The red box in (B) indicates the reverse complement sequence of the T7 promoter. C. Schematic illustrating the results in (A-B). Transcription using a template with a single T7 promoter results in the production of both sense and antisense transcripts, which differ in length by the size of the T7 promoter. Solid and dotted lines indicate DNA and RNA, respectively. D. Native PAGE analysis of T7 transcripts generated using DNA template with a single T7 promoter (1), DNA template without the T7 promoter (2), and gel-purified 512B ssRNA as a template (3). RNA template alone (4) was compared with (3).

It was also reported that T7 pol can generate RNA using another RNA as a template (17,18). To examine this possibility and to determine exactly how the dsRNA byproduct was made, we performed 3’ - and 5’ -RACE analyses of the RNA complementary to 512B (c512B) (Figure 2B).

The results showed that c512B starts primarily from the 3’ end of 512B (Figure 2C). Note that the template sequence in this end has little resemblance to the canonical T7 promoter. 3’ -RACE of the c512B further revealed that c512B ends with the sequence complementary to the T7 promoter. These results suggest that the c512B is produced by transcription in reverse orientation from the promoter-less DNA end of the 512B template, rather than by RNA-dependent RNA polymerization. To further confirm that c512B is synthesized by promote-independent, DNA-dependent transcription, we performed the T7 transcription using 512B RNA as a template and 512B DNA template without the T7 promoter at either end. Consistent with the notion that c512B is generated from the promoter-independent, DNA-dependent RNA polymerization, dsRNA was generated from the DNA even without the T7 promoter, but not from the RNA template (Figure 2D). The transcription reaction using 512B RNA template did not produce any detectable level of dsRNA or transcription product aside from the input RNA.

### Promoter-independent antisense transcription occurs for a wide range of DNA with various end sequences and structures

Our finding that promoter-independent T7 pol transcription begins at the promoter-less DNA end raised the question of whether this activity depends on the DNA end sequence or structure. Note that the four RNAs tested above (512A-D) have different sequences but share the common 6 nucleotides at the promoter-less end of the DNA template (Table S1). The templates were also commonly prepared by PCR amplification, and thus are expected to harbor blunt ends. To examine the importance of the DNA end sequence and structure, we compared three sequence variants that differ from the original 512B template near the promoter-less end (i.e. transcriptional start site for c512B) (Figure 3A). In the varied end sequence, we embedded unique restriction sites (XmaI, BamHI and KpnI), which enabled us to generate ends containing 5’− and 3’−overhangs by restriction digestion. The Klenow reaction was also performed to ensure that the original 512B template harbors blunt ends.

**Figure 3.**
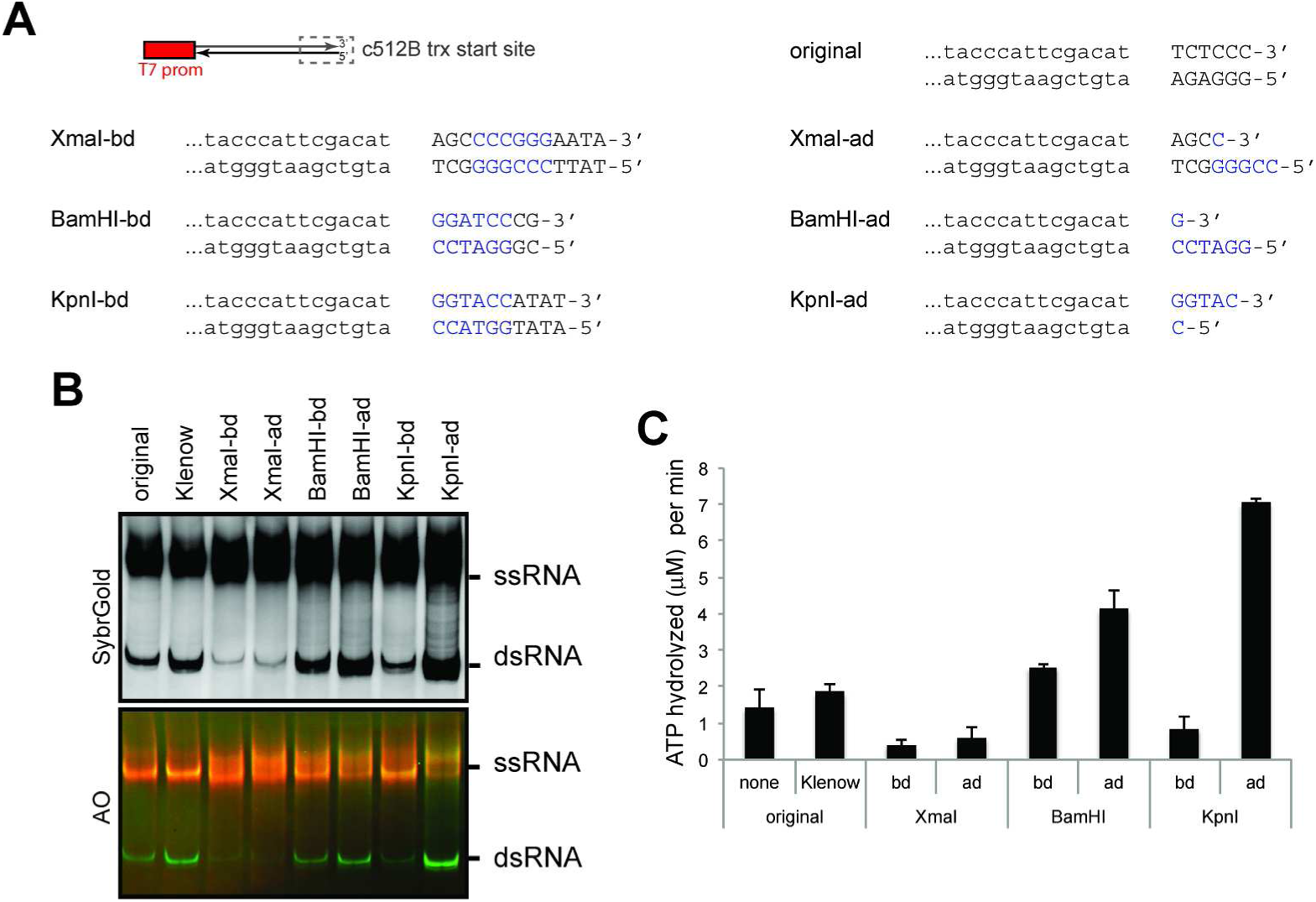
Impact of the DNA termini sequence and structure on the promoter-independent antisense transcription. A. Sequence and structure variants of DNA templates tested in this study. All have a single T7 promoter for transcription of 512B. The sequence near the promoter-less end (*i.e.* transcriptional start site for the antisense transcript, c512B) were varied and shown in capital letters. DNA end structural variants were generated by restriction digestion of the listed DNA. Restriction sites are colored blue, and −bd and −ad indicate before and after digestion, respectively. B. Native PAGE analysis of the T7 transcripts produced using the original 512B template before and after the Klenow reaction, and its sequence variants before and after respective restriction digestions. C. MDA5 ATPase assay of the T7 transcripts in (B).

Comparison of the transcripts from these DNA templates before and after Klenow or restriction digestion suggests that the end sequence and structures can influence the efficiency of promoter-independent antisense transcription, as measured by the dsRNA byproduct on the native gel (Figure 3B) and MDA5 ATPase activity (Figure 3C). Note that the XmaI sequence lowered the level of dsRNA, whereas restriction digestion increased the level of dsRNA regardless of the sequence or overhang type. In particular, digestion with KpnI, which generates a 3’ overhang, increased the level of dsRNA most significantly. While the higher level of dsRNA with KpnI-digested DNA is consistent with the previous report that a 3’ overhang promotes antisense transcription (19), our finding suggests that even DNA with a 5’ overhang (BamHI-ad) or a blunt end (Klenow) have a significant level of antisense transcription. These results thus suggest that the promoter-independent antisense transcription is a more general phenomenon than was previously thought, and can be influenced, but not eliminated by both DNA end sequence and structure.

### Modified nucleotides affect the level of dsRNA byproduct, not its MDA5-stimluatory activity

It has been shown that co-transcriptional incorporation of modified nucleotides can decrease to a certain extent the immunogenicity of T7 transcripts (4,14,20,21). These modified nucleotides include pseudouridine (Ψ), 1-methylpseudouridine (m1Ψ), □−methyladenosine (m6A), and 5-methylcytosine (m5C) (Figure 4A). We asked whether the presence of modified nucleotides suppress the formation of dsRNA byproducts or immune-stimulatory activity of the dsRNA. To test these hypotheses, we performed four independent transcription reactions, in each of which Ψ, m1Ψ, m6A or m5C completely replaced U, U, A or C, respectively. Intriguingly, Ψ, m1Ψ and m5C, but not m6A, suppressed dsRNA byproduct formation, as measured by native gel analysis (Figure 4B). The cellular MDA5 signaling activity and MDA5 ATPase activity also correlated with the level of dsRNA byproduct observed in the transcripts (Figures 4C & 4D).

**Figure 4.**
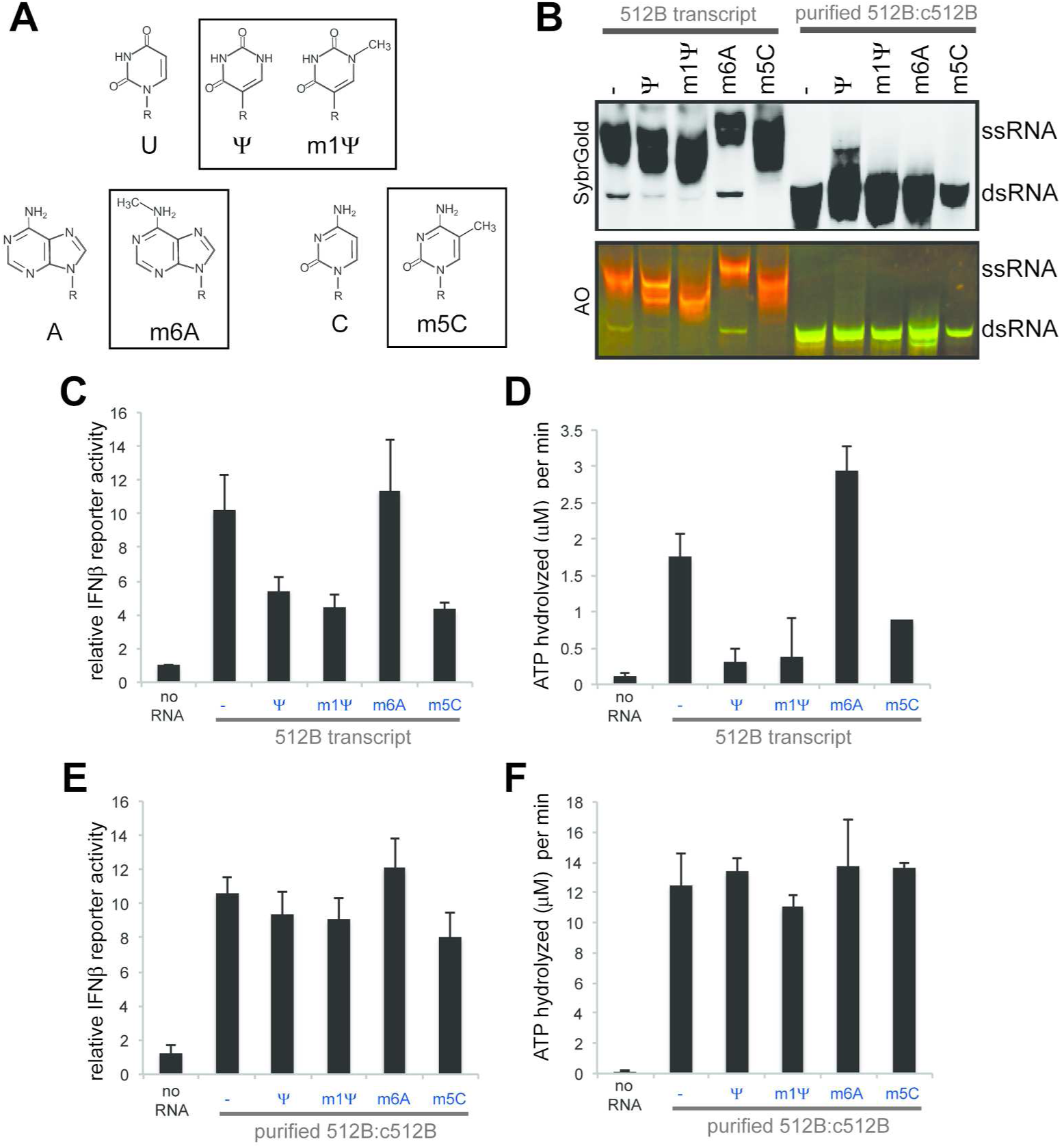
Modified nucleotides affect the level of dsRNA byproduct, not its MDA5− stimluatory activity. A. Chemical structures of the modified nucleotides used in this study (boxed) and their respective unmodified counterparts. B. Native PAGE analysis of the T7 transcripts for 512B or the dsRNA species (512B:c512B) prepared with or without the indicated modified nucleotides. The dsRNA was prepared by co-transcription of the two RNA strands using two independent templates and was purified using RNase I_f_, which selectively degrades residual ssRNA while maintaining dsRNA. Modified nucleotides were used in place of the respective unmodified nucleotide. C. MDA5-stimulatory activity of 512B transcripts in (B), as measured by IFNβ promoter-driven dual luciferase assay. D. ATPase activity of MDA5 with 512B transcripts in (B). E. MDA5-stimulatory activity of purified 512B:c512B dsRNA in (B), as measured by IFNβ promoter-driven dual luciferase assay. F. ATPase activity of MDA5 with purified 512B:c512B dsRNA in (B).

We next asked whether incorporation of Ψ, m1Ψ, m6A or m5C has any effect on the dsRNA’s ability to stimulate MDA5, in addition to the level of dsRNA byproduct formation. To address this question, we examined MDA5 signaling and ATPase activities using modified and unmodified dsRNA of 512B:c512B hybrids. The dsRNA was prepared by co-transcription of the two RNA strands using two independent templates and was purified using RNase I_f_, which selectively degrades residual ssRNA while maintaining dsRNA (Figure 4B). The results show that Ψ, m1Ψ, m6A or m5C had little impact on the MDA5-stimulatory activity of the dsRNA (Figures 4E & 4F). It is noteworthy that another nucleotide modification, adenosine deamination (i.e. inosine) by the cellular enzyme ADAR1, significantly suppresses the MDA5 activity (22–24). Given that inosine weakens Watson-Crick base-pairs, while Ψ, m1Ψ, m6A and m5C do not, this result is consistent with the notion that MDA5 primarily recognizes dsRNA backbone structure with little specific contacts with RNA bases (25). Thus, the observed impact of Ψ, m1Ψ and m5C in lowering the immunogenicity of T7 transcript is at least in part due to their abilities to lower the dsRNA byproduct formation by T7 pol, rather than by decreasing the MDA5-stimulatory activity of the dsRNA byproduct.

### Low concentrations of Mg can suppress the promoter-independent antisense transcription

The above result suggested that the transcription reaction conditions can affect the dsRNA byproduct formation. We thus performed a more systematic analysis of the reaction condition using varying concentrations of individual reaction components. These include T7 pol, DNA template, NTP and MgCl_2_, of which a range of concentrations have been used in literature. We also tested an increasing concentration of NaCl, because non-specific DNA binding by T7 pol was previously shown to be suppressed in buffers with high ionic strength (26). By both in-house produced T7 pol (hT7 pol) and commercial T7 pol (cT7 pol), we observed little dependence of the dsRNA byproduct formation on the concentration of T7 pol (Figure 5A). Concentrations of the template, NTP and NaCl also had a minimal impact on the level of dsRNA byproduct formation (Figures 5B–D). Intriguingly, decreasing the concentration of MgCl_2_ from 30 mM to 5 mM significantly reduced dsRNA byproduct formation, to the point that dsRNA was barely detectable at 5 mM MgCl_2_ (Figure 5E). Note that the total yield of RNA was not significantly affected by the reduction in MgCl_2_. This was unexpected because it is often considered necessary to use MgCl_2_ in excess over NTP (which is 20 mM in total). The observed impact of MgCl_2_ on dsRNA production was similarly observed with the MDA5 ATPase assay, which showed that decreasing concentration of MgCl_2_ reduced the level of the ATPase activity (Figure 5F).

**Figure 5.**
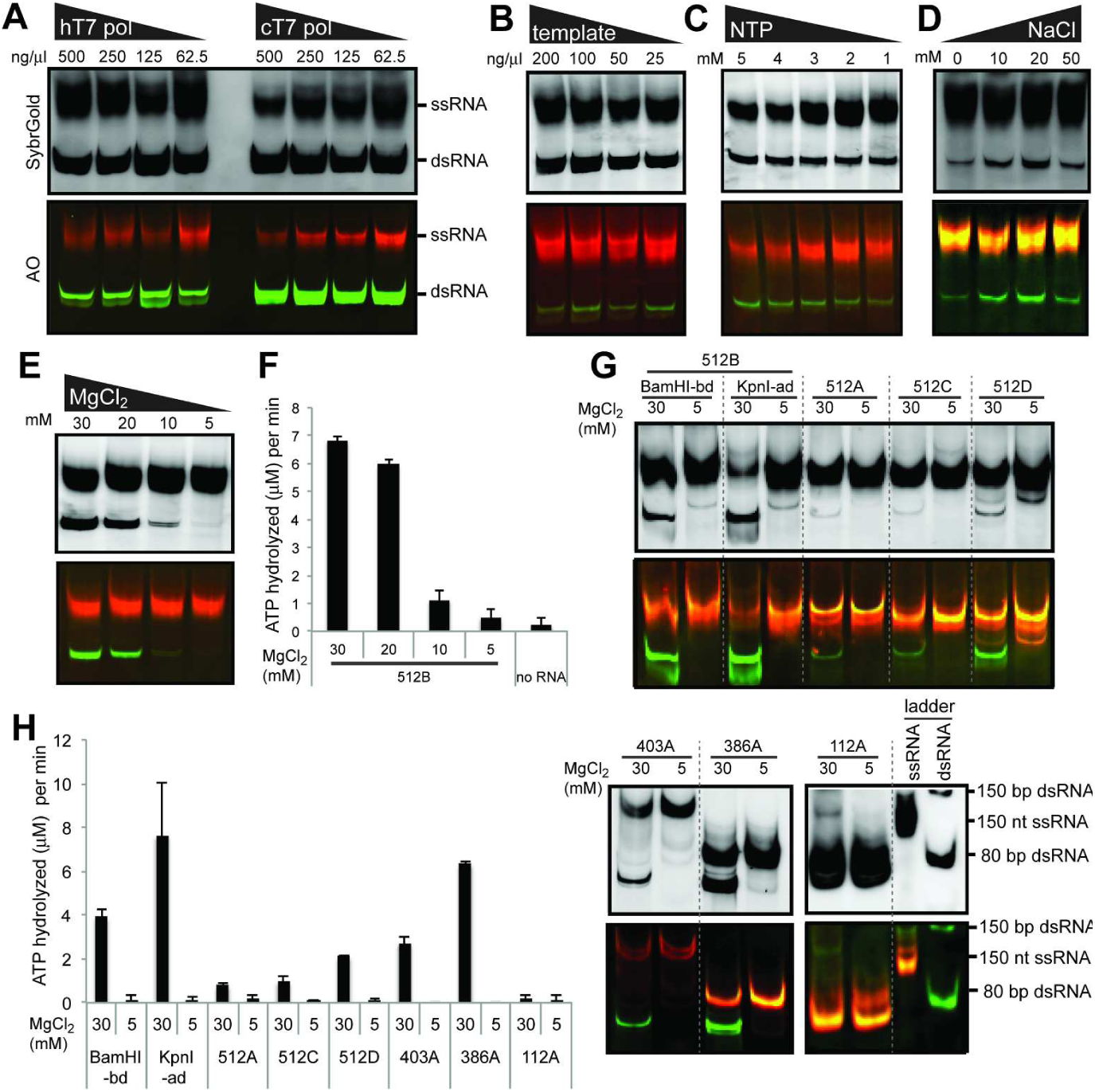
Impact of the transcription reaction conditions on dsRNA byproduct formation. A. Native PAGE analysis of the T7 transcripts for 512B prepared using a decreasing concentration of T7 pol. In-house T7 pol (hT7 pol) and commercial T7 pol (cT7 pol, ThermoFisher) were compared. B-E. Native PAGE analysis of the T7 transcripts for 512B prepared using a decreasing concentration of the DNA template (B), NTP (C) and MgCl_2_ (E), and an increasing concentration of NaCl (D). F. MDA5 ATPase analysis of the T7 transcripts in (E). G. Native PAGE analysis of the T7 transcripts for 512B variants (BamHI-bd and KpnI-ad), 512A, 512C, 512D, 403A, 386A and 112A generated using MgCl_2_ of 30 mM and 5 mM. See Table S1 for the template sequence. Note that for shorter RNAs (~150 nt), dsRNA migrates more slowly than ssRNA of an equivalent size. H. MDA5 ATPase analysis of the T7 transcripts in (G).

To examine whether the observed impact of MgCl_2_ on dsRNA byproduct formation is generally applicable, we examined other templates with varying DNA sequences, structure and lengths. In all cases, including the one with a 3’ overhang (KpnI-ad), we found that reducing MgCl_2_ from 30 mM to 5 mM significantly lowered the level of dsRNA byproduct, as measured by both native gel analysis and MDA5 ATPase assay (Figures 5G & 5H). Although additional ssRNA bands appear for some RNAs at low MgCl_2_ (Figure 5G), the overall lack of dsRNA byproduct suggests that lowering the MgCl_2_ concentration can be used as a general method to suppress the aberrant dsRNA byproduct formation.

## DISCUSSION

It has been known that T7 *in vitro* transcripts are often immunogenic. However, the precise origin of the immunogenicity−i.e. the identity of the RNA species and its immune-stimulatory mechanism−has been unclear. This study provides an answer to these questions by showing that T7 pol often produces long dsRNA byproducts that can robustly activate the cytosolic sensors RIG-I and MDA5 in 5’ppp-dependent and -independent manners. The dsRNA is formed by hybridization of the intended sense transcript and its fully complementary antisense transcript. The antisense RNA is produced by promoter-independent transcriptional initiation from a DNA end, which differs from previously reported erroneous behaviors of T7 pol, such as 3’ -extension of RNA, RNA-dependent RNA polymerization and DNA-primed RNA synthesis (16–18,27). Our finding also shows that this promoter-independent transcription is observed with a wide range of DNA templates, and is not limited to the DNA end with 3’ overhangs as previously reported (19). Thus, our work highlights yet another level of complexity of T7 pol activity, and has broad implications for studies utilizing T7-transcribed RNAs.

Our work also offers novel methods to quantitatively detect such dsRNA byproduct formation. While previous studies often utilized dsRNA-specific antibodies, such as J2, for detecting dsRNA, such methods cannot distinguish between secondary structures within ssRNA vs. long dsRNA byproducts formed by sense-antisense hybridization. Instead, we propose the MDA5 filament formation and ATPase assays as robust and convenient methods to quantitatively measure dsRNA contaminants, either in T7 pol transcripts or in any other RNA products. In particular, MDA5 filament analysis by EM allows one to examine the extent of the continuous duplex length, thereby unambiguously distinguishing between intrinsic secondary structures within ssRNA and fully duplexed RNA byproducts. Of note, our work also demands re-evaluation of the conventional method of determining RNA purity, *i.e.* denaturing gel electrophoresis. Considering that intended sense and antisense transcripts differ by ~20 nt (the size of the T7 promoter, Figure 2), denaturing gel would be inefficient in separating sense and antisense strands for long RNA transcripts. Instead we here used native gel electrophoresis, which allowed us to separate ssRNA and dsRNA products and to distinguish them using acridine orange staining. While native gel analysis alone can be complicated for RNAs with intrinsic secondary structures, we believe that a combination of native gel electrophoresis, MDA5 filament formation and ATPase assays would enable a comprehensive analysis of RNA transcript of interest.

Finally, our study provides previously unappreciated methods to suppress the dsRNA byproduct formation. These include using modified nucleotides and reducing MgCl_2_ concentration, as well as altering template sequence in the promoter-less terminus. Unlike the conventional notion that RNA containing modified nucleotides are less immunogenic, we found that modified nucleotides in fact suppress the T7 pol’s aberrant activity of generating immunogenic dsRNA byproduct, rather than decreasing the MDA5-stimulatory activity of dsRNA. We also found that Mg, among all variables tested, unexpectedly plays the most significant role in regulating dsRNA byproduct synthesis. Intriguingly, a previous study showed that HPLC purification can deplete immune-stimulatory dsRNA (5). Although the exact identity of the dsRNA byproduct in this study is not clear, an optimal transcription reaction that minimizes such byproduct formation would help in improving the purity of RNA, and could potentially alleviate the need for the HPLC purification. Altogether, the current work provides novel insights into the previously enigmatic properties of the T7 pol–an origin, detection and regulation of the immunogenic byproduct.

## FUNDING

SA was funded by Cancer Research Institute. This work was supported by NIH grants (R01AI106912, R01AI111784 and R21AI130791) to SH.

## Conflict of interest

None.

## Materials and Methods

### Plasmids and Proteins

Mammalian expression plasmids for RIG-I and MDA5 and the bacterial expression plasmid for MDA5∆N (residue 287-1025) were described previously (15). Protocols to express and purify MDA5∆N was reported in (25). Briefly, the protein was expressed in BL21(DE3) at 20˚C for 16-20 hr following induction with 0.5 mM IPTG. Cells were lysed by high pressure homogenization using an Emulsiflex C3 (Avestin), and the protein was purified by a combination of Ni-NTA and heparin affinity chromatography and size exclusion chromatography (SEC) in 20 mM Hepes, pH 7.5, 150 mM NaCl and 2 mM DTT. T7 polymerase was expressed in BL21(DE3) using the plasmid pT7-911Q. The T7 pol protein was purified by Ni-NTA followed by size exclusion chromatography (SEC) in 50 mM Tris, pH 7.5, 100 mM NaCl and 1 mM EDTA, and stored in 50% glycerol.

### T7 transcription

Sequences for the DNA templates are shown in Table S1. Unless mentioned otherwise, templates for *in vitro* transcription were prepared by PCR reactions. Transcription was performed in the reaction containing 250 mM Hepes (pH 7.5), 30 mM MgCl_2_, 2 mM Spermidine, 40 mM DTT, 0.1 mg/ml BSA, 5 mM NTP (5 mM each), 0.5 mg/ml T7 pol and 0.25 mg/ml DNA template. Reaction was incubated at 37˚C for 4 hr and DNA template was digested with DNase I. Trasnscript was purified by Phenol:chloroform extraction, ethanol precipitation and using the QIAquick PCR purification kit (Qiagen). Transcripts (160 ng each) were analyzed by 1X TBE 6% polyacrylamide gel. RNAs were visualized by SybrGold staining (Thermofisher) or by acridine orange (AO, in 25 μg/ml) staining (Sigma Aldrich). Fluorescent gel images were obtained using the FLA9000 gel scanner (GE Healthcare). For the AO fluorescent images, ssRNA-specific images were obtained using the 473 nm and 792 nm filters for excitation and emission, respectively. For dsRNA-specific fluorescent images, the 532 nm and 554 nm filters were used for excitation and emission, respectively.

### 5’ - and 3’ - RACE

For 5’ -RACE, transcripts were subject to reverse transcription using a primer specific to the RNA of interest. Prior to RT, the primer and RNA were incubated at 95˚C for 5 min in the presence of 2 mM EDTA, and cooled down to 4˚C for 2 min. High Capacity cDNA reverse transcription kit (Applied Biosystems) was used according to the manufacturer’s protocol. After RT, RNA was removed with RNase A, and cDNA was purified using QIAquick PCR purification kit. The 3’ end of the first strand cDNA was extended with poly A tails using the Terminal dNTP transferase (NEB). The second strand cDNA was synthesized by using the oligo d(T)-anchor primer. For 3’ -RACE, 3’ end of the RNA was first extended with poly A tails using the poly A polymerase (NEB). The first strand and second strand cDNAs were synthesized using the oligo d(T)-anchor primer and a primer bearing the internal sequence of the RNA of interest. For both 5’ – and 3’ -RACE, the final product was PCR amplified and subjected to Sanger sequencing (Quitara Biosciences). Primers used in the analysis are shown in Table S2.

**Table S2.**
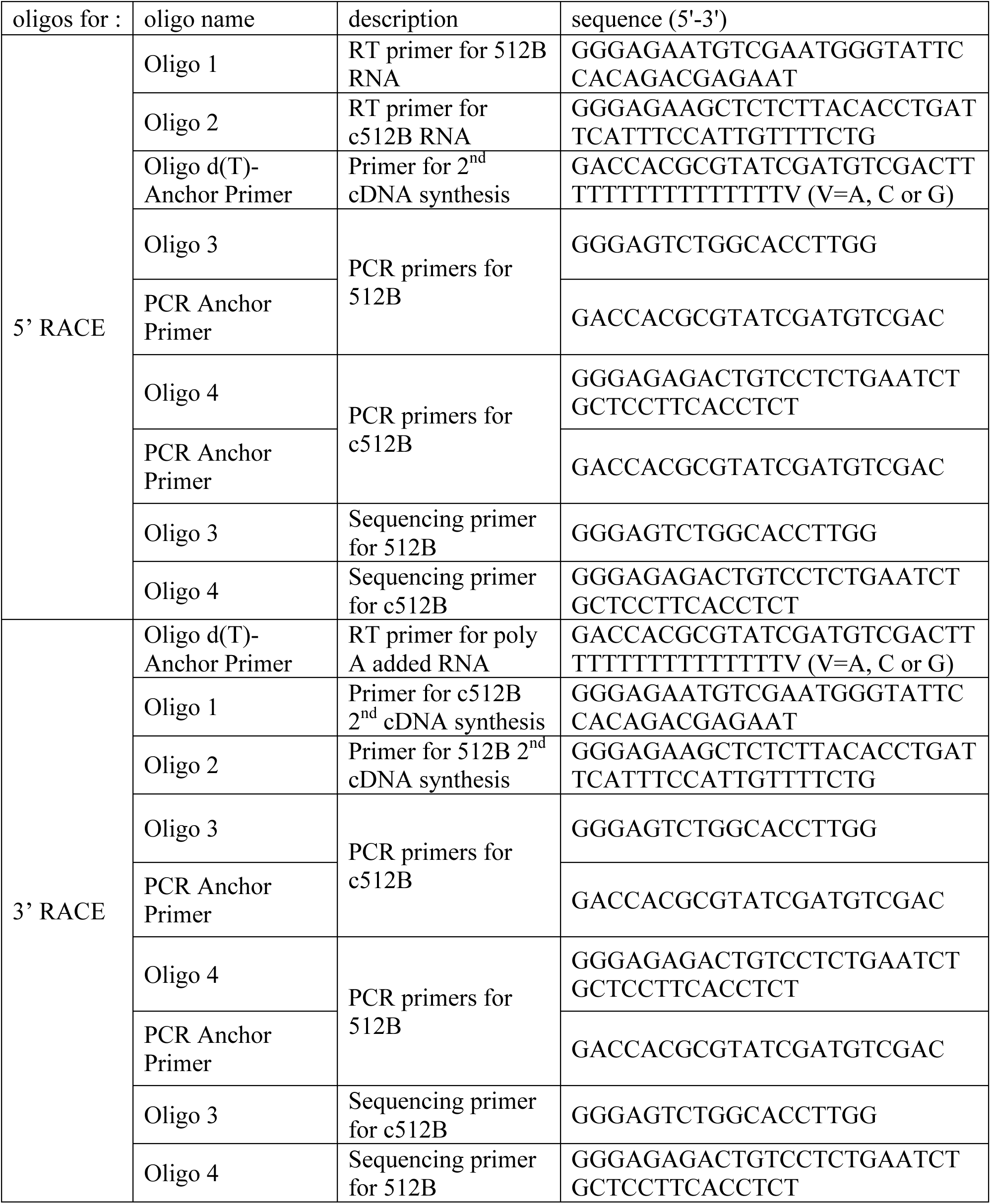
Sequences of DNA oligonucleotides for RACE.

### IFNβ promoter-driven dual luciferase assay

293T cells were maintained in 48-well plates in Dulbecco’s modified Eagle medium (Cellgro) supplemented with 10% heat-inactivated fetal calf serum and 1% penicillin/streptomycin. At ~90% confluence, cells were transfected with the pFLAG-CMV4 plasmids encoding RIG-I (10 ng) or MDA5 (5 ng), the IFNβ- promoter driven firefly luciferase reporter plasmid (100 ng) and a constitutively expressed Renilla luciferase reporter plasmid (pRL-CMV, 10 ng) using lipofectamine2000 (Life) according to the manufacturer’s protocol. The medium was changed 6-8 hr after the first transfection and the cells were additionally transfected with in vitro transcribed RNA (0.5 µg) or high molecular weight polyIC (0.5 µg, Invivogen). Cells were lysed ~20 hr post-stimulation and IFNβ promoter activity was measured using the Dual Luciferase Reporter assay (Promega) and a Synergy2 plate reader (BioTek). Firefly luciferase activity was normalized against Renilla luciferase activity.

### ATPase assay

The ATP hydrolysis activity of MDA5 was measured using Green Reagent (Enzo Life Sciences). MDA5ΔN (500 nM) was pre-incubated with RNA (2 ng/µl) in buffer A (20 mM HEPES pH 7.5, 150 mM NaCl, 1.5 mM MgCl_2_ and 2 mM DTT), and the reaction was initiated by addition of 2 mM ATP at 37°C. Aliquots (10 µl) were withdrawn before and 15 min after ATP addition, and were quenched with 100 mM EDTA on ice. The Green Reagent (90 µl) was added to the quenched reaction at a ratio of 9:1, and OD_650_ was measured using a Synergy2 plate reader (BioTek).

### Electron Microscopy

MDA5ΔN (450 nM) was incubated with RNA (2 ng/µl) in buffer A for 10 min at RT followed by addition of 1 mM ADP•AlF_x_ on ice. ADP•AlF_x_ was prepared by mixing ADP, AlCl_3_ and NaF in a molar ratio of 1:1:3. Prepared filaments were adsorbed to carbon-coated grids (Ted Pella) and stained with 0.75% uranyl formate as described(28). Images were collected using a Tecnai G^2^ Spirit BioTWIN transmission electron microscope at 49,000x magnification.

### RNase III / RNase I_f_ digestion

100 ng/µl of RNA was incubated with indicated concentrations of RNase III (NEB) or RNase I_f_ (NEB) in 10 mM Tris, pH 8.5 and 2 mM MgCl_2_ at 37°C for 30 min. The reaction was terminated using 1/10 volume of proteinase K (NEB) and incubated at 25°C for 20 min. RNA was purified with Direct-zol RNA MiniPrep Kit (Zymo research) followed by QIAquick PCR purification kit (Qiagen).

